# Emotion-induced frontal *α* asymmetry predicts relapse after discontinuation of antidepressant medication

**DOI:** 10.1101/2023.07.05.547831

**Authors:** Isabel M. Berwian, Marius Tröndle, Carlota de Miquel, Anastasios Ziogas, Gabor Stefanics, Henrik Walter, Klaas Enno Stephan, Quentin J.M. Huys

## Abstract

**Background:** One in three patients relapse after antidepressant discontinuation. Thus, the prevention of relapse after achieving remission is an important component in the long-term management of Major Depressive Disorder (MDD). However, no clinical or other predictors are established. Frontal reactivity to sad mood as measured by fMRI has been reported to relate to relapse independently of antidepressant discontinuation and is an interesting candidate predictor.

**Methods:** Patients (n=56) who had remitted from a depressive episode while taking antidepressants underwent EEG recording during a sad mood induction procedure prior to gradually discontinuing their medication. Relapse was assessed over a six-months follow-up period. 35 healthy controls were also tested. Current source density of the EEG power in the *α* band (8-13Hz) was extracted and alpha-asymmetry was computed by comparing the power across two hemispheres at frontal electrodes (F5 and F6).

**Outcomes:** Sad mood induction was robust across all groups. Reactivity of *α*-asymmetry to sad mood did not distinguish healthy controls from patients with remitted MDD on medication. However, the 14 (25%) patients who relapsed during the follow-up period after discontinuing medication showed significantly reduced reactivity in *α*-asymmetry compared to patients who remained well. This EEG signal provided predictive power (69% out-of-sample balanced accuracy).

**Interpretation:** A simple EEG-based measure of emotional reactivity may have clinical utility in the management of antidepressant discontinuation.

**Funding:** Swiss National Science Foundation project grant 320030L_153449 / 1 to QJMH, Stiftung Deutsche Depressionshilfe to HW and QJMH, a Deutsche Forschungsgemeinschaft (DFG) grant (WA 1539/5-1) to HW, EMDO Stiftung to QJMH and the René and Susanne Braginsky Foundation and Clinical Research Priority Programme “Molecular Imaging” at the University of Zurich to KES.

## 1 Introduction

Depression is the leading cause of disability and amongst the most burdensome disorders worldwide^1^. The reason for this is in no small part due to its chronicity. The time spent in depression rises linearly with age amongst those affected^2^, effectively taking away a substantial part of an individual’s life. The risk of an episode is initially strongly affected by stressful life events, but over time becomes a self-propelling force^3;4^, with the risk of chronicity increasing with each episode^5^. Hence, the goal of depression treatment must not only be the resolution of a particular episode, but also the long-term prevention of relapses.

The choice of whether to discontinue an antidepressant medication plays an important role in this setting because relapses frequently follow discontinuation: around one third of those who discontinue antidepressant medication suffer a relapse within months of discontinuation^6;7;8;9;10^. Unfortunately, clinically salient features have little value in predicting who will relapse after discontinuation^11;12^. For instance, estimates of the impact of the number of previous episodes diverge diametrically^6;8^, while length of remission on medication does not relate to relapse risk^6;7;8;13;14^.

As such, it may be useful and necessary to examine more affective, cognitive or neurobiological measures as predictors of relapse risk after antidepressant discontinuation^15^. Current mood, as well as mood reactivity, but not cognitive reactivity have been shown to be associated with relapse independent of antidepressant discontinuation^16;17;18^. Sadness-induced maladaptive cognitions^19;20^ and functional magnetic resonance imaging (fMRI) measures of induced sadness^21^ or self-blame^22^ have all been reported to predict relapse. Hence, the induction of emotions associated with the depressed state appears to be a promising probe. Farb and colleages^21^ further found that brooding related to the sadness-induced fMRI measure and subsequent relapse. They^23^ further replicated and extended their neuroimaging results in a randomized controlled trial showing that somatosensory deactivation is associated with subsequent relapse, while a reduction of left dorso-lateral prefrontal cortex activity due to prophylactic treatment might protect from later relapse. The procedure by Farb *et al*.^21^,^23^ is clinically particularly appealing as it involves simply presenting participants with sad and neutral movies during neuroimaging. However, the reliance on fMRI is a hindrance to clinical translation due to availability, costs and contraindications to MRI.

In an attempt to maximise the potential for clinical applicability, we recorded brain activity with EEG during sad mood induction. An EEG marker that has been related to both depressive episodes and emotion induction is frontal *α* assymmetry (FAA). FAA power is defined as the contrast of alpha power over the frontal right vs frontal left electrodes. Throughout this paper, FAA will refer to the asymmetry in *α* power such that negative values indicate more left alpha power and positive values more right FAA power. Note that *α* power (frequency range of 8 to 13 Hz) is inversely related to cortical activity as measured by PET and functional MRI recordings^24;25;26^, such that more left FAA power is indicative of more right cortical activity^27^. FAA power appears to be driven by frontal areas^28^ similar to those predictive of relapse in fMRI measurements^21^.

Interpretation of FAA is often based on the approach-withdrawal model suggesting that more left FAA power relates to withdrawal motivation, while more right FAA power relates to approach motivation. According to this model, an individual’s FAA reflects their affective style across contexts^29;30^ and might be a trait marker of current and past depression^31^ (see Thibodeau *et al*.^32^ for a review). Recent meta-analyses based on more coherent and rigorous inclusion and quality criteria, however, found no consistent evidence for FAA as a diagnostic marker^33^. The relationship between FAA and depression might be moderated by covariates such as comorbid anxiety and gender^32;34;33;35^. FAA in combination with covariates might also be a promising prognostic marker, as older women who subsequently responded to SSRIs showed more right FAA^36^ and FAA was associated with the onset of first episodes^37^.

The capability model^38^ suggests that it is the FAA specifically during emotional challenges which might be the more reliable indicator of affective style and motivational differences as it assesses the abilities to respond to, or more specifically regulate, emotions. Indeed, stronger left shifts of FAA power during emotion induction differentiate better between currently depressed, previously depressed, and healthy participants than static rest FAA measures^39;40;41^. Here, we further enhanced the reliability by applying a current source density transformation to the data^41^. Similarly, there is evidence for a left-laterization of FAA power during emotion induction in children at risk for depression^42^ and participants with dysphoria^43^ (see^44^ for a review).

The specific role of responses to mood induction in the setting of antidepressant discontinuation as opposed to more general relapse prediction has not been addressed. In addition, relapse specifically after antidepressant discontinuation may involve both pharmacological but also psychological mechanisms. The knowledge that a potentially effective medication is being withdrawn may have a nocebo effect, and this may impact realistic clinical settings^45^.

In this study, we therefore examined if neural responses to mood induction assessed with EEG have potential in predicting relapse after open-label, naturalistic antidepressant discontinuation. Patients with remitted Major Depressive Disorder (MDD) were followed up after discontinuing their antidepressant medication and after viewing sad and neutral movies while EEG was recorded.

Based on previous results and in an attempt to extend the results by Farb and colleagues to EEG^21^, we aimed to examine the role and predictive power of sadness-induced FAA and rumination in relation to disease and medication state and relapse after antidepressant discontinuation. Of note, we have previously reported that clinical and demographic variables do not predict relapse in this sample^12^, highlighting the importance of identifying other potentially predictive measurements.

## 2 Methods

### 2.1 Participants

The AIDA study^12^ was a bicentric study with sites in Zurich, Switzerland and Berlin, Germany. It recruited healthy controls and patients with either severe or recurrent DSM-IV-TR Major Depressive Disorder (MDD)^46^, who had remitted from a Major Depressive Episode while taking antidepressant medication (i.e. they did not fulfill the criteria of a depressive episode at study entry and had a score of less than 7 on the Hamilton Depression Rating Scale), and who were intending to discontinue their medication (see Supplementary Methods S1.1 full in- and exclusion criteria). Participation in the EEG recording was an additional voluntary component of the study at the Zurich site only. All participants gave written, informed consent, received monetary compensation for their time, and indicated whether they were willing to participate in an additional EEG session, which lasted approximately three hours. The study was approved by the Cantonal Ethics Commission Zurich (BASEC: PB_2016-0.01032; KEK-ZH: 2014-0355).

### 2.2 Study design

After providing consent, participants underwent a baseline session to assess inclusion criteria by trained staff, acquire baseline self-report and demographic characteristics, and plan the study assessments and discontinuation (see Berwian *et al*.^12^ for full details). They then underwent the EEG session prior to gradually tapering their antidepressant medication over up to 18 weeks and entering a 6-months follow-up. Participants were contacted by telephone at weeks 1, 2, 4, 6, 8, 12, 16 and 21 after the end of medication discontinuation to assess relapse, and were encouraged to contact the study team in case of a worsening of the mental state. If the telephone assessment indicated a possible relapse, they were invited for an in-person structured clinical interview. Relapse was defined as fulfilling the criteria for a Major Depressive Episode according to DSM-IV-TR. A final assessment was conducted if relapse occurred or 6 months after completing discontinuation. (MDD)^46^. Study team members were not involved in treatment decisions. These remained fully with the treating physician and the participants. All participants additionally underwent two main assessment sessions involving fMRI, behavioural and biological testing (see Berwian *et al*.^12^,^47^,^48^).

### 2.3 EEG experiment

Prior to the EEG recordings, participants were asked to abstain from alcohol and coffee for 24h. A 64-channel EasyCap (EASYCAP GmbH, Herrsching, Germany) adapted to head-size was fitted and Abralyt gel applied. The reference electrode was placed on the nose tip. Additional channels were placed underneath the left eye for electrooculogram and on the back to acquire electrocardiogram. We aimed for impedances *<*5kΩ and accepted impedances *<*10kΩ.

Prior to the experiment, participants rated one 45s extract from each of four movies (The Champ, Terms of Endearment, Stepmom and The Sixth Sense; Farb *et al*.^21^) in terms of their subjective sadness on 7-point Likert scales. The two movies with the highest subjective sadness were then chosen as the sad movies, each lasting approximately 3 minutes. Two neutral movies of 2 min 15 second duration were also chosen as control stimuli. All movie clips were then divided into segments lasting approximately 45 seconds, i.e. the sad movies were divided into four, and the neutral movies into three 45s clips. During the experiment, participants were comfortably seated in a shielded room and monitored via video. They passively viewed the clips in a fixed order: neutral movie 1 (3 clips) - sad movie 1 (4 clips) - neutral movie 2 (3 clips) - sad movie 2 (4 clips). After each 45s clip, they rated their subjective sadness on the same 7-point Likert scale.

### 2.4 Self-and Observer-rated measurements

In- and exclusion criteria and disease history and course were assessed with the German translations of the Structured Clinical Interview for DSM-IV (SCID) I and II^49^. Residual symptoms of depression were assessed with the Hamilton Depression Rating Scale^50^ and the Inventory of Depressive Symptomatology^51^ while residual anxiety was measured with the GAD-7^52^. Rumination^53;54^ was also assessed. Finally, we assessed verbal intelligence with the Mehrfachwahl Wortschatz Test^55^, working memory with the digit span backwards test from the Wechsler Adult Intelligence Scale^56^, cognitive processing speed with the Trail Making Test A^57^ and executive processing speed with the Trail Making Test B^57^.

### 2.5 Analysis

Data analysis was performed using Matlab 9.4.0.813654 (www.mathworks.com) with SPM 12 v7219 (https://www.fil.ion.ucl.ac.uk/spm/), MARA v1.2 (https://github.com/irenne/MARA)^59^, EEGLab 14.1.1b (https://sccn.ucsd.edu/eeglab/index.php) and CSD^60^ (http://psychophysiology.cpmc.columbia.edu/Soft-ware/CSDtoolbox/) toolboxes. Data were bandpass filtered between 0.5 and 40 Hz and divided into 1s epochs with 500ms overlap. Artifact were removed first by using ICA and MARA^59^, and epochs rejected by thresholding. As recommended^61^, Current Source Density was then estimated using the CSD toolbox^60^. CSD is the Laplacian, i.e. the second spatial derivative, of the scalp surface voltage, and is used to estimate the current projected towards the skull and hence giving rise to the observed scalp potentials. CSD minimizes the influence of volume conduction in the measured signal, and therefore enhances the detection of distinct focal patterns in the distribution of alpha power over the scalp. In line with previous literature on FAA in MDD, the power in the *α* band was estimated as average power in 8-13 Hz applying a fast Fourier transform to the CSD estimates. Finally, the data was transformed with a logarithm to the base of 10 and multiplied by 10.

A priori analyses focused on the contrast between electrodes F5 and F6^28^. FAA was defined as right minus left frontal activity (F6-F5) of the log-transformed data. eFAA was defined as FAA during while watching sad movies minus FAA while watching neutral movies. Thus, negative eFAA indicate stronger induction of left lateralization during the emotional challenge. Unless indicated otherwise, group comparisons were two-sample two-tailed t-tests. If a Kolmogorov-Smirnov indicated that the data was not normally distributed, non-parametric tests were applied instead. Individual correlations between eFAA and self-reported sadness were performed by averaging the eFAA over each of the 14 45s clips performing a linear correlation with the self-reported sadness after each of the clips. Logistic regressions were performed using the Matlab function glmfit.m. We included residual symptoms to control for baseline severity and brooding due its relation to sadness-induced BOLD activity and relapse in the study by Farb et al.^21^ as covariates. As we had already established that all available standard clinical and demographic variables do not predict relapse in this study, we did not include them in the present analyses^12^. To assess out-of-sample prediction, leave-one-out cross-validation was performed, i.e. we first estimated beta coefficients for a logistic regression using all but one subject and then used these estimates and a fixed threshold of 0.5 to predict relapse for the left-out subject. The significance of the balanced accuracy was assessed using a binomial test comparing with 0.5. We only included residual symptoms, brooding and eFAA in this analyses due to the reasons outlined above. No critical hyperparameters were tuned and, thus, nested cross-validation was not required.

Medication load was computed as 100 × *d/m/w*, with *d* being the dose in mg, *m* being the maximal allowed dose in mg in the Swiss compendium (www.compendium.ch), and *w* being the weight.

## 3 Results

### 3.1 Participants

123 patients and 66 healthy controls were included overall, and 56 of the 85 patients and 35 of the 40 healthy controls at the EEG site (Zurich) agreed to participate in the additional EEG study. Of the 56 patients, 48 (86%) reached a study endpoint (Fig. 1), with 14 (29%) suffering a relapse. Patients and controls did not differ on age, sex, BMI and neuropsychological variables (Tab. 1). Patients who would go on to relapse and those who would remain well after discontinuation did not differ in terms of number of prior episodes, onset age, medication load, and residual depression and anxiety symptoms as well as demographic and neuropsychological variables prior to antidepressant discontinuation (Tab. 1).

**Table 1:** Sample characteristics.

|  | Controls | Patients | *p* | Relapsers | Non-relapsers | *p* |
| --- | --- | --- | --- | --- | --- | --- |
| N | 35 | 56 |  | 14 | 34 |  |
| Gender (Women) | 25 | 44 | 0.96 | 11 | 26 | 1 |
| Handedness (left) | 0 | 2 | 0.87 | 1 | 1 | 0.98 |
| Age | 32.97±10.61 | 34.36±11.01 | 0.56 | 35.14±9.11 | 33.12±11.69 | 0.57 |
| BMI | 22.92±4.03 | 23.76±4.15 | 0.35 | 22.52±3.17 | 23.81±4.01 | 0.29 |
| HAM-D<sub>17</sub> | 0.55±1.03 | 1.60±1.66 | **0.0019** | 1.38±1.04 | 1.42±1.64 | 0.94 |
| IDS-C | 0.71±1.19 | 3.32±2.81 | **4.4x10<sup>-6</sup>** | 2.15±1.28 | 3.30±2.80 | 0.16 |
| GAD-7 | 1.88±1.98 | 2.80±2.00 | **0.038** | 2.71±1.64 | 2.59±2.08 | 0.84 |
| Onset age | – | 23.77±8.98 | – | 23.29±6.78 | 23.24±9.21 | 0.99 |
| # prior episodes | – | 2.70±1.31 | – | 2.64±1.22 | 2.65±1.32 | 0.99 |
| Medication Load | – | 0.81±0.42 | – | 0.70±0.35 | 0.83±0.45 | 0.35 |
| MWTB | 28.03±4.36 | 28.05±4.36 | 0.98 | 29.21±3.75 | 27.26±4.83 | 0.18 |
| TMT A | 23.85±5.91 | 24.16±6.47 | 0.82 | 21.49±5.48 | 24.43±6.82 | 0.16 |
| TMT B | 59.25±22.47 | 59.12±16.57 | 0.98 | 53.79±14.04 | 61.64±18.31 | 0.16 |
| Digit Span | 8.00±3.70 | 7.14±2.18 | 0.17 | 6.79±1.67 | 7.03±2.24 | 0.72 |

Shown are mean ± 1 st. dev; *p*-values show $\chi^2$ for categorical and otherwise 2-sample *t*-tests. BMI = Body Mass Index; Digit span = digit span backwards from Wechsler Adult Intelligence Scale<sup>56</sup>; GAD-7 = 7-item Generalized Anxiety Disorder Scale<sup>58</sup>; HAM-D<sub>17</sub> = Hamilton Scale for Depression, 17-item version<sup>50</sup>, MWTB = Mehrfachwahl Wortschatz Test<sup>55</sup>; TMT = Trail Making Test<sup>57</sup>;

**Figure 1:**
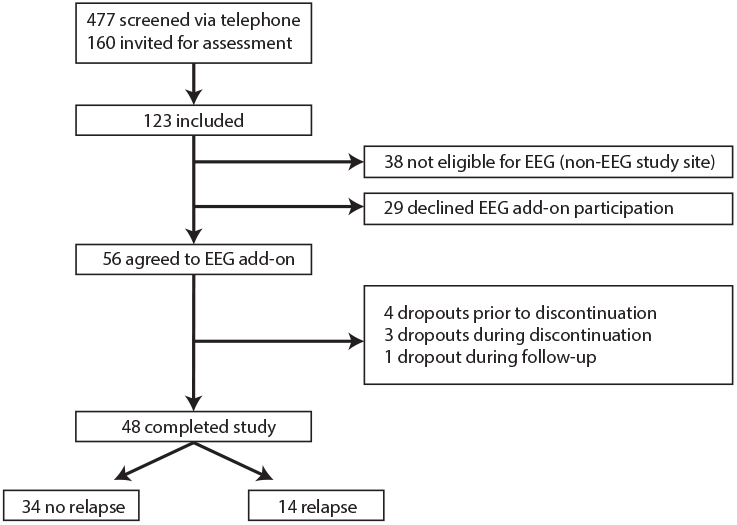
Consort diagram.

### 3.2 Sadness-evoked *α* power

The sadness induction procedure proved effective. All groups showed a highly significant (all p < 0.0001) but modest (around 1.5 points on a 7-point Likert scale) increase in sadness during the sad compared to the neutral movies (Fig. 2A) that did not differ between any two groups (all p>.4).

**Figure 2:**
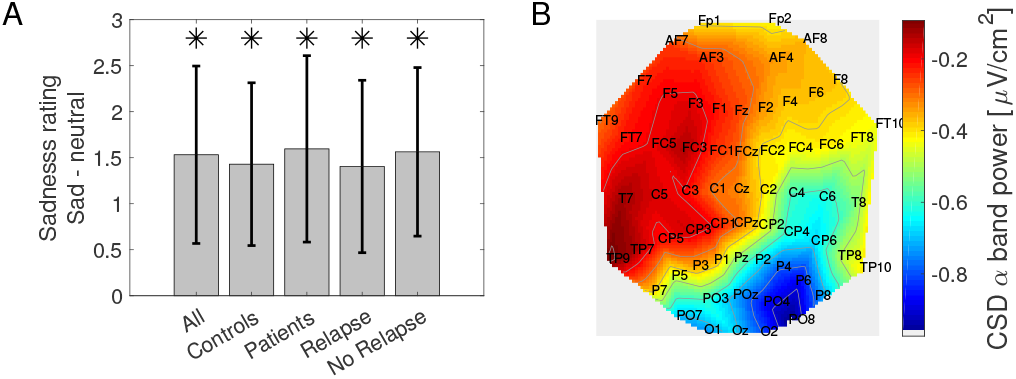
**A**: Difference in sadness ratings between sad and neutral movies for all groups and subgroups. Error bars show 1 st. dev. * indicates p < .0001 difference from zero. **B**: Scalp plot of current source density (CSD) estimates in the *α* band comparing sad to neutral movie viewing, showing an emotion-induced shift in *α* power towards the left across all participants in the study.

Non-parametric Wilcoxon tests were used as a Kolmogorov-Smirnov test indicated that eFAA and the correlation of eFAA and sadness were not normally distributed (all p < 0.001). Note that t-tests led to a similar pattern of results as reported below. Across all groups, we observed an emotion-induced FAA (eFAA), i.e. the sadness manipulation induced a shift in *α* power in current source density between electrodes F5/F6 (Z=-3.5, p<0.001). The topography of the overall distribution of the effect (Fig. 2B) indicated a broad shift in *α* power towards the left, but all statistical analyses focus on the electrode pair F5/F6 as pre-specified. The shift was present at a trend level in controls and more clearly in the patient group as a whole (Z=-1.85, p=0.06 and Z=-3.00, p=0.003), respectively; Fig 3B). Patients and controls did not differ overall (Z=0.49, p=0.63). However, amongst the patients, the effect was driven by those who would go on to remain well after antidepressant discontinuation (Fig. 3C; Z=-4.01, p<0.001) and not present in those who went on to relapse (Fig. 3C; p=0.35), resulting in a significant difference (Z=2.82, p=0.005) with an effect size of Cohen’s d’=1.01.

**Figure 3:**
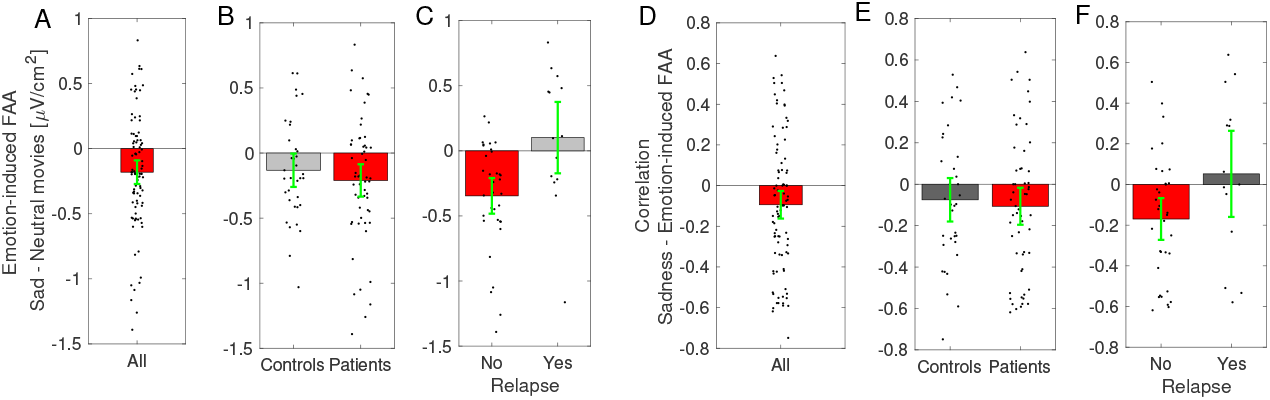
*α* lateralization induced by sadness. Negative values indicate a lateralization to the left. **A-C**: Shift in *α* power towards the left induced by sad movies compared to neutral movies. **D-F**: Correlation of emotion-induced shift in *α* power with self-reported sadness induced by movies. Green error bars show 95% confidence intervals.

Furthermore, this shift in power was more pronounced the larger the self-reported sadness induced by the movies. This was true when examining all subjects overall (Fig. 3D, Z=-2.71, p=0.007), and amongst patients (Fig. 3E, right, Z=-2.34, p=0.02) but not amongst controls (Fig. 3E, left, Z=-1.40, p=0.16), though the difference between patients and controls was not significant (Z=0.40, p=0.68). Amongst patients, on the other hand, there was a trend-level difference between those who did and did not go on to relapse (Z=1.86, p=0.06, Fig. 3F), with a correlation present amongst those who did not relapse (Z=-2.09, p=0.004) but not amongst those who went on to relapse (p=0.64).

### 3.3 Correlation with rumination and related measures

As a consequence of the relapse/no relapse effect, emotion-induced *α* asymmetry also correlated with the depression severity at study endpoint (Fig. 4, r=0.39, p=0.0049). The direction of this effect suggests that patients who showed a an attenuated response in terms of left lateralization of FAA due to the emotion induction, i.e. in line with emotional insensitivity, were more likely to have symptoms at study endpoint. Notably, eFAA did not correlate with residual depression symptoms at baseline (r=-0.038, p=0.72).

**Figure 4:**
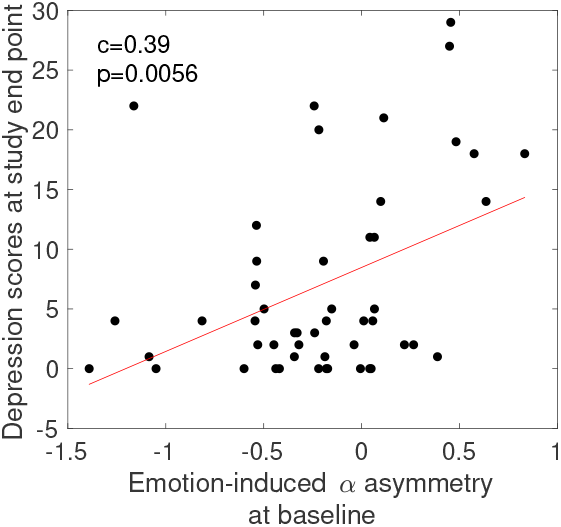
Emotion-induced *α* asymmetry correlated with the Hamilton score at study endpoint.

Farb *et al*.^21^ reported that midfrontal sadness-induced BOLD reactivity was correlated with rumination scores and indeed that this BOLD signal mediated the relationship between relapse and rumination. In our sample, there was no correlation between sadness-induced *α* asymmetry and either total rumination scores, or the brooding or reflection aspects of rumination.

### 3.4 Prediction

Next, we asked whether the EEG measure could improve relapse prediction above and beyond the residual depression symptoms and brooding. This logistic regression achieved an area under the curve (AUC) of 0.88 (Fig. 5A), and a balanced accuracy with the optimally selected threshold of 0.86 (Fig. 5B). Exclusion of eFAA resulted in a worse performance with an AUC of 0.72 (Fig. 5A).

**Figure 5:**
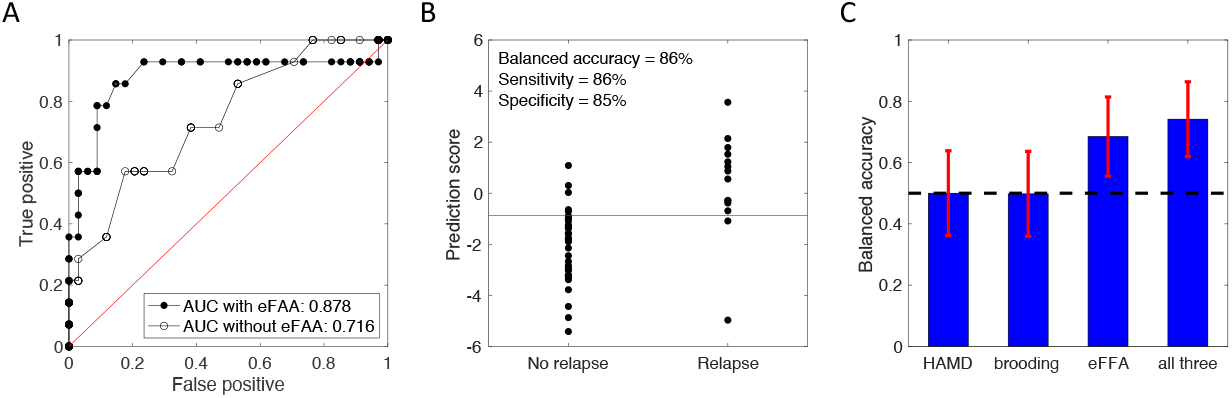
Relapse prediction. **A**: The area under the curve is increased from 0.72 without including eFAA (empty circles) to 0.88 with eFAA (black dots). **B**: Optimal threshold showing classification on the weighted prediction score including eFAA. **C**: Out-of-sample prediction. eFFA predicts relapse well above chance (0.69), while depression symptoms and brooding do not. The black dashed line indicates chance level. Error bars indicate 95% Bayesian credible intervals. eFFA = emotion-induced frontal *α* asymmetry; HAMD = Hamilton 17 rating scale for depression.

However, the above results do not represent true ‘prediction’ as the parameters are fitted to the dataset that the model is then evaluated on. We next asked whether eFAA could improve out-of-sample prediction above and beyond the residual symptoms and brooding. We estimated likely out-of-sample prediction through a leave-one-out cross-validation procedure. While residual depression symptoms and brooding did not predict relapse (balances accuracy=0.5, p=0.44 and p=0.56, respectively), eFFA predicted relapse with balanced accuracy of 0.69 (p=0.007). Combining the three measures improved the balanced accuracy to 0.74 (p=0.0004; Fig. 5C).

### 3.5 Exploratory analyses

FAA during the neutral or sad condition alone did not differ between groups (see Fig. S1 and Fig. S2) and did not predict subsequent relapse out-of-sample (p=0.55 and p=0.76, respectively). To ensure that our analyses are robust with regard to data transformation and electrode pairs, we replicated all analyses using the contrast F4/F3, an electrode pair often used in the literature^33^, a data transformation with the natural log for F6/F5 and data transformation according to F6-F5/F6+F5. All analyses led to the same pattern of results as reported in this paper with similar levels of prediction for F4/F3 with an out-of sample accuracy of 0.70 (p=0.003) for eFAA. Data transformation with the natural log led to identical prediction levels for the F6/F5 contrast as reported above, while data transformation according to (F6-F5)/(F6+F5) led to slightly weaker prediction results with an out-of-sample accuracy of 0.61 (p=0.06) for eFAA.

## 4 Discussion

Relapses after discontinuing antidepressant medications are common, and occur in around one third of cases within 6 months irrespective of the previous duration of medication^6;7;8^. Our findings are very much in keeping with this. In the larger study, of which this is a subset, we have recently found that standard clinical variables have no predictive power^12^. This highlights the need for other predictive markers, e.g. based on neurocognitive processes, to guide clinical decision-making.

Here, we examined a relatively simple measure that could credibly be translated to clinical situations: watching emotional movie clips during EEG recording prior to initiating the medication discontinuation. The sad movies resulted in an emotion-induced frontal *α* asymmetry (eFAA), i.e. it induced a small but significant self-reported increase in sadness, and this was accompanied by a shift in frontal *α* current density from the right to the left, suggesting relative left hypoactivation of the frontal cortex. This effect robustly differentiated patients who would go on to relapse from those who would remain well after discontinuing antidepressants, with no significant difference between the group of remitted patients as a whole and healthy never-depressed controls.

Several points are worth noting. First, FAA induced by sad vs neutral movies was not able to distinguish between patients and controls. Although some studies were able to differentiate between these two groups by using this marker (e.g. Stewart *et al*.^39^ ; Grünewald *et al*.^62^), others failed to do so and recent meta-analyses failed to find a significant difference on FAA between depressed participants and healthy controls^33;63^. This conflicting evidence could be due to a variety of reasons. In their meta-analysis, Thibodeau *et al*.^32^ concluded that several variables moderate the effect sizes of FAA as a marker for depression status. Larger effects were observed in adult samples with shorter EEG recording periods, with Cz references and mid-frontal recording sites, and in younger childhood depression samples. Medication has also been previously found to reduce distinctions between patient and control groups^64^, raising the possibility that the antidepressant medication normalizes frontal asymmetry. Of note, our entire patient sample was on antidepressant medication during the EEG assessment which could explain the lack of a group difference. In addition, our findings are in line with evidence that FAA relates to the subsequent course of depression^36;37^.

Second, the BOLD findings by Farb *et al*.^21^ and Farb *et al*.^23^ suggested that an increase in prefrontal reactivity to sad emotions and a lack of a reduction of this increase due to prophylactic treatment might index a higher relapse risk, respectively. Here, we observe a reduction in or no shift in FAA induced by the sad emotion in patients who went on to relapse. While this is neutral with respect to the overall activation, it does suggest a reduction in neural reactivity to the emotional material rather than an increase. Furthermore, our finding indicates that patients who relapse are more akin to the controls than those who show a resilient course. This raises the possibility that those individuals who have maintained neural markers of emotional reactivity while on antidepressants have had to acquire emotion regulation to maintain an euthymic mood, while those who have a blunted emotion reactivity on antidepressants suffer a relapse because they have not acquired these mood regulation skills prior to discontinuing the medication. Our findings are consistent with reports that better immediate response to a cognitive restructuring intervention correlated with more alpha power in F3 vs F4^65^.

Depression is also often characterized by a blunted response to negative stimuli^66;67^ and such a blunted response measured in terms of late positive potentials has been associated with worse outcome after Cognitive Behavioral Therapy^68^ and an increase in dysphoria^69^. Blunted responses to negative stimuli might be marker for a group of patients with a subtype of depression with worse prognosis. Hence, an alternative explanation for our pattern of results is that patients who are unresponsive to negative stimuli are more likely to relapse. Both explanations are in line with the result that less left lateralization in FAA during the emotional challenge was followed by more symptoms of depression at study endpoint.

Third, FAA has recently been related to left-lateral and dorsolateral prefrontal sources in depression^28^ that appear more lateral than the medial prefrontal sources identified using BOLD^21^. However, the replication by Farb *et al*.^23^ in a larger randomized controlled trial indicated that a lack of attenuation of activity in the dorso-lateral prefrontal cortex due to prophylactic cognitive therapy with a well-being focus or mindfulness based cognitive therapy was related to subsequent relapse risk. Hence, the regions identified by Farb *et al*.^23^ seem to overlap with the areas generating FAA in the sample by Smith *et al*.^28^. This pattern of results indicates that the regions driving the BOLD effects by Farb *et al*.^23^ may be similar to those that drive the FAA effects in our sample.

Fourth, technical aspects of the associations and prediction results are worth reiterating. When fitting a model to the entire dataset, we found an association between residual symptoms and brooding and relapse even without eFAA, but that it was increased by eFAA. However, these analyses are not true prediction in the sense that they do not reflect prediction on unseen data. To approach this issue, we iteratively put the data of one subject aside, trained the model on all other data, and then asked how well the association generalized to the data not seen during training. This identified a predictive power with an out-of-sample balanced accuracy of 74% when eFFA was included, while residual symptoms and brooding alone predicted relapse at chance level (50%). This suggests that 4 individuals need to be tested with EEG to predict outcome of one more patient correctly than when using simpler clinical procedures.

Our study has a number of strengths, but also limitations. The most important strength and limitation is the naturalistic design. Because discontinuation was open-label, all of the current results could be driven either by pharmacological or psychological (nocebo) effects. Disentangling pharmacological and psychological effects of discontinuation requires a placebo-controlled study. While the inability to make causal statements is clearly a limitation, the naturalistic design also strengthens the clinical validity of the study as it captures relapse in the relevant setting where both pharmacological and psychological forces are jointly at play. Second, while we have used sadness as the target emotion, our study does not speak to whether this emotion is strictly necessary, and whether other emotions, or only aspects of emotions such as arousal, valence or regulation strategy^70^ are relevant. The third limitation is the sample size, which is modest. However, we do not the challenge in acquiring such samples. While quantitative EEG markers have long been thought to be promising, the literature suffers from heterogeneity, there is evidence for a substantial publication bias, and the sensitivity and specificity remain insufficient for clinical applications at present^71;72^. The current results (particularly in terms of prediction) should hence motivate further studies, but must be interpreted with caution given these caveats. The sample is representative of the Swiss population and only includes white participants. The results might not generalize to members of other ethnicities.

In conclusion, we have examined a simple EEG procedure in individuals who seek to discontinue their antidepressant medication after having achieved stable remission. If the current findings replicate, they may serve as the basis of a simple, potentially portable solution to support clinical decision-making at the time of antidepressant discontinuation.

## Data access

Sharing data of this study conforms to EU regulations (GDPR) and Swiss data protection regulations. The decision will be based on the acceptance by the study team that a valid and timely scientific question, based on a written protocol, has been posed by those seeking to access the data. Safeguarding of ethical standards and legal obligations will be required. The role of the original study team will need to be acknowledged. Please contact the corresponding author via email to request access to the data. Data access for questions of scientific integrity may additionally be regulated via the funder.

## Code access

Access to code will be provided upon request.

## 4.1 Acknowledgements

We thank Tania Villar for support with data acquisition, and Inga Schnürer for support in running the study.

## 4.3 Contributions

QJMH and HW conceived and designed the study with input from KES. IMB, CM, MT, GS and AZ collected the data under the supervision of QJMH. KES was the study sponsor in Zurich. QJMH, KES and HW acquired funding for the study. MT, IMB, AZ, GS and QJMH planned and performed the analyses. QJMH completed the analyses and wrote the manuscript. All authors provided critical comments, read and approved the manuscript.

## 5 Supplementary Material

### S1 Supplementary Methods

#### S1.1 In- and Exclusion Criteria

Participants fulfilling the following inclusion criteria were eligible for participation in the study:

1. age 18-55 years
2. ability to consent and adhere to the study protocol
3. written informed consent
4. fluent in written and spoken German.

Patients had to additionally fulfil the following criteria:

1. currently under medical care with a psychiatrist or general practitioner for remitted Major Depressive Disorder and willing to remain in care for the duration of the study (approx. 9 months)
2. informed choice to discontinue medication (including willingness to taper the medication over at most 12 weeks) that was independent of study participation
3. clinical remission (HAMD_17_ of less than 7) had been achieved under therapy with Antidepressant Medication (ADM) without having undergone manualized psychotherapy; with no other concurrent psychotropic medication and had been maintained for a minimum of 30 days,
4. consent to information exchange between treating physician and study team members regarding inclusion/exclusion criteria and past medical history.

Any of the following exclusion criteria led to exclusion of an participant. This included the following general criteria

1. any disease of type and severity sufficient to influence the planned measurement or to interfere with the parameters of interest (This includes neurological, endocrinological, oncological comorbidities, a history of traumatic or other brain injury, neurosurgery or longer loss of consciousness.)
2. premenstrual syndrome (ICD-10 N94.3).

and MRI-related criteria

1. MRI-incompatible metal parts in the body,
2. inability to sit or lie still for a longer period,
3. possibility of presence of any metal fragments in the body,
4. pregnancy,
5. pacemaker, neurostimulator or any other head or heart implants,
6. claustrophobia and
7. dependence on hearing aid.

For patients the following additional criteria would led to exclusion:

1. current psychotropic medication other than antidepressants,
2. questionable history of major depressive episodes without complicating factors,
3. current acute suicidality,
4. lifetime or current axis II diagnosis of borderline or antisocial personality disorder,
5. lifetime or current psychotic disorder of any kind, bipolar disorder,
6. current posttraumatic stress disorder, obsessive compulsive disorder, or eating disorder
7. current drug use disorder (with the exception of nicotine) or within the past 5 years.

Healthy controls were excluded if there was a lifetime history of Diagnostic and Statistical Manual of Mental Disorders 4th ed., text rev.; DSM-IV-TR;^73^ axis I or axis II disorder with the exception of nicotine dependence.

### S2 Supplementary Results

FAA while participants watched neutral or sad movies did not differentiate between patients and controls or relapsers and non-relapsers as evident in Fig. S1 and Fig. S2.

**Figure S1:**
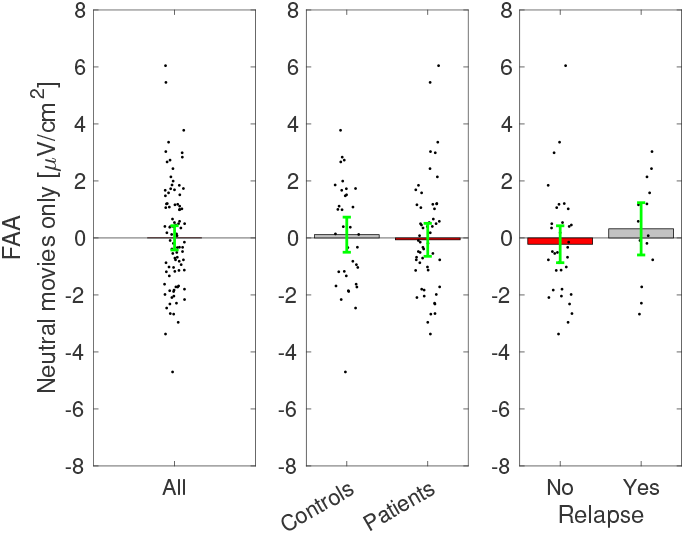
*α* lateralization during neutral movies for all participant groups. Negative values indicate a lateralization to the left. Green error bars show 95% confidence intervals.

**Figure S2:**
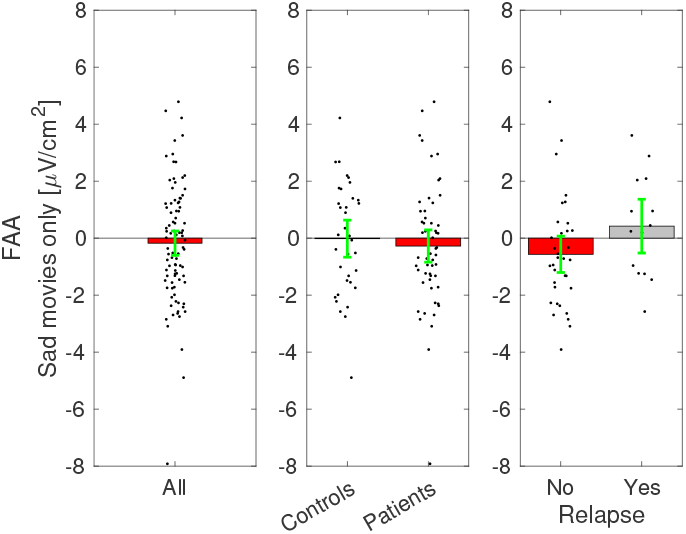
*α* lateralization during sad movies for all participant groups. Negative values indicate a lateralization to the left. Green error bars show 95% confidence intervals.

## References

[1] WHO. Depression and Other Common Mental Disorders: Global Health Estimates. Geneva, CH (2017).

[2] Angst, J., Gamma, A., Sellaro, R., Lavori, P.W. & Zhang, H. Recurrence of bipolar disorders and major depression. a life-long perspective. Eur Arch Psychiatry Clin Neurosci 253, 236–240 (2003).

[3] Angst, J. Epidemiology of depression. Psychopharmacology (Berl) 106 Suppl, S71–S74 (1992).

[4] Kendler, K.S., Thornton, L.M. & Gardner, C.O. Stressful life events and previous episodes in the etiology of major depression in women: An evaluation of the “kindling” hypothesis. Am. J. Psychiatry 157, 1243–51 (2000).

[5] Hollon, S.D., Shelton, R.C., Wisniewski, S., Warden, D., Biggs, M.M. et al. Presenting characteristics of depressed outpatients as a function of recurrence: preliminary findings from the star*d clinical trial. J Psychiatr Res 40, 59–69 (2006).

[6] Viguera, A.C., Baldessarini, R.J. & Friedberg, J. Discontinuing antidepressant treatment in major depression. Harv Rev Psychiatry 5, 293–306 (1998).

[7] Geddes, J.R., Carney, S.M., Davies, C., Furukawa, T.A., Kupfer, D.J. et al. Relapse prevention with antidepressant drug treatment in depressive disorders: a systematic review. Lancet 361, 653–661 (2003).

[8] Kaymaz, N., van Os, J., Loonen, A.J.M. & Nolen, W.A. Evidence that patients with single versus recurrent depressive episodes are differentially sensitive to treatment discontinuation: a meta-analysis of placebo-controlled randomized trials. J Clin Psychiatry 69, 1423–1436 (2008).

[9] Borges, S., Chen, Y.F., Laughren, T.P., Temple, R., Patel, H.D. et al. Review of maintenance trials for major depressive disorder: a 25-year perspective from the us food and drug administration. The Journal of clinical psychiatry 75, 205–214 (2014).

[10] Lewis, G., Marston, L., Duffy, L., Freemantle, N., Gilbody, S. et al. Maintenance or discontinuation of antidepressants in primary care. The New England journal of medicine 385, 1257–1267 (2021).

[11] Berwian, I.M., Walter, H., Seifritz, E. & Huys, Q.J.M. Predicting relapse after antidepressant withdrawal - a systematic review. Psychol Med 47, 426–437 (2017).

[12] Berwian, I.M., Wenzel, J.G., Kuehn, L., Schnuerer, I., Seifritz, E. et al. Low predictive power of clinical features for relapse prediction after antidepressant discontinuation in a naturalistic setting. Sci Rep 12, 11171 (2022).

[13] Glue, P., Donovan, M.R., Kolluri, S. & Emir, B. Meta-analysis of relapse prevention antidepressant trials in depressive disorders. Aust N Z J Psychiatry 44, 697–705 (2010).

[14] Andrews, P.W., Kornstein, S.G., Halberstadt, L.J., Gardner, C.O. & Neale, M.C. Blue again: perturbational effects of antidepressants suggest monoaminergic homeostasis in major depression. Front Psychol 2, 159 (2011).

[15] Buckman, J.E.J., Underwood, A., Clarke, K., Saunders, R., Hollon, S.D. et al. Risk factors for relapse and recurrence of depression in adults and how they operate: A four-phase systematic review and meta-synthesis. Clinical psychology review 64, 13–38 (2018).

[16] van Rijsbergen, G.D., Bockting, C.L., Berking, M., Koeter, M.W. & Schene, A.H. Can a one-item mood scale do the trick? predicting relapse over 5.5-years in recurrent depression. PLoS One 7, e46796 (2012).

[17] van Rijsbergen, G.D., Bockting, C.L., Burger, H., Spinhoven, P., Koeter, M.W. et al. Mood reactivity rather than cognitive reactivity is predictive of depressive relapse: a randomized study with 5.5-year follow-up. Journal of Consulting and Clinical Psychology 81, 508 (2013).

[18] van Rijsbergen, G.D., Burger, H., Hollon, S.D., Elgersma, H.J., Kok, G.D. et al. How do you feel? detection of recurrent major depressive disorder using a single-item screening tool. Psychiatry research 220, 287–293 (2014).

[19] Segal, Z.V., Kennedy, S., Gemar, M., Hood, K., Pedersen, R. et al. Cognitive reactivity to sad mood provocation and the prediction of depressive relapse. Arch Gen Psychiatry 63, 749–755 (2006).

[20] Segal, Z.V., Bieling, P., Young, T., MacQueen, G., Cooke, R. et al. Antidepressant monotherapy vs sequential pharmacotherapy and mindfulness-based cognitive therapy, or placebo, for relapse prophylaxis in recurrent depression. Arch Gen Psychiatry 67, 1256–1264 (2010).

[21] Farb, N.A.S., Anderson, A.K., Bloch, R.T. & Segal, Z.V. Mood-linked responses in medial prefrontal cortex predict relapse in patients with recurrent unipolar depression. Biol Psychiatry 70, 366–372 (2011).

[22] Lythe, K.E., Moll, J., Gethin, J.A., Workman, C.I., Green, S. et al. Self-blame-selective hyperconnectivity between anterior temporal and subgenual cortices and prediction of recurrent depressive episodes. JAMA Psychiatry 72, 1119–1126 (2015).

[23] Farb, N.A.S., Desormeau, P., Anderson, A.K. & Segal, Z.V. Static and treatment-responsive brain biomarkers of depression relapse vulnerability following prophylactic psychotherapy: Evidence from a randomized control trial. Neuroimage Clin 34, 102969 (2022).

[24] Larson, C.L., Davidson, R.J., Abercrombie, H.C., Ward, R.T., Schaefer, S.M. et al. Relations between pet-derived measures of thalamic glucose metabolism and eeg alpha power. Psychophysiology 35, 162–169 (1998).

[25] Oakes, T.R., Pizzagalli, D.A., Hendrick, A.M., Horras, K.A., Larson, C.L. et al. Functional coupling of simultaneous electrical and metabolic activity in the human brain. Human brain mapping 21, 257–270 (2004).

[26] Laufs, H., Holt, J.L., Elfont, R., Krams, M., Paul, J.S. et al. Where the bold signal goes when alpha eeg leaves. NeuroImage 31, 1408–1418 (2006).

[27] Allen, J.J.B., Coan, J.A. & Nazarian, M. Issues and assumptions on the road from raw signals to metrics of frontal eeg asymmetry in emotion. Biol Psychol 67, 183–218 (2004).

[28] Smith, E.E., Cavanagh, J.F. & Allen, J.J.B. Intracranial source activity (eloreta) related to scalp-level asymmetry scores and depression status. Psychophysiology 55 (2018).

[29] Davidson, R.J. Emotion and affective style: Hemispheric substrates (1992).

[30] Davidson, R.J. Affective style and affective disorders: Perspectives from affective neuroscience. Cognition & emotion 12, 307–330 (1998).

[31] Henriques, J.B. & Davidson, R.J. Left frontal hypoactivation in depression. J Abnorm Psychol 100, 535–545 (1991).

[32] Thibodeau, R., Jorgensen, R.S. & Kim, S. Depression, anxiety, and resting frontal eeg asymmetry: a meta-analytic review. Journal of abnormal psychology 115, 715–729 (2006).

[33] van der Vinne, N., Vollebregt, M.A., van Putten, M.J.A.M. & Arns, M. Frontal alpha asymmetry as a diagnostic marker in depression: Fact or fiction? a meta-analysis. NeuroImage. Clinical 16, 79–87 (2017).

[34] Bruder, G.E., Stewart, J.W. & McGrath, P.J. Right brain, left brain in depressive disorders: Clinical and theoretical implications of behavioral, electrophysiological and neuroimaging findings. Neuroscience and biobehavioral reviews 78, 178–191 (2017).

[35] Nusslock, R., Shackman, A.J., McMenamin, B.W., Greischar, L.L., Davidson, R.J. et al. Comorbid anxiety moderates the relationship between depression history and prefrontal eeg asymmetry. Psychophysiology 55 (2018).

[36] Arns, M., Bruder, G., Hegerl, U., Spooner, C., Palmer, D.M. et al. Eeg alpha asymmetry as a gender-specific predictor of outcome to acute treatment with different antidepressant medications in the randomized ispot-d study. Clin Neurophysiol (2015).

[37] Nusslock, R., Shackman, A.J., Harmon-Jones, E., Alloy, L.B., Coan, J.A. et al. Cognitive vulnerability and frontal brain asymmetry: common predictors of first prospective depressive episode. Journal of abnormal psychology 120, 497 (2011).

[38] Coan, J.A., Allen, J.J.B. & McKnight, P.E. A capability model of individual differences in frontal eeg asymmetry. Biol Psychol 72, 198–207 (2006).

[39] Stewart, J.L., Bismark, A.W., Towers, D.N., Coan, J.A. & Allen, J.J.B. Resting frontal eeg asymmetry as an endophenotype for depression risk: sex-specific patterns of frontal brain asymmetry. Journal of abnormal psychology 119, 502–512 (2010).

[40] Stewart, J.L., Coan, J.A., Towers, D.N. & Allen, J.J.B. Frontal eeg asymmetry during emotional challenge differentiates individuals with and without lifetime major depressive disorder. Journal of affective disorders 129, 167–174 (2011).

[41] Stewart, J.L., Coan, J.A., Towers, D.N. & Allen, J.J.B. Resting and task-elicited prefrontal eeg alpha asymmetry in depression: support for the capability model. Psychophysiology 51, 446–455 (2014).

[42] Lopez-Duran, N.L., Nusslock, R., George, C. & Kovacs, M. Frontal eeg asymmetry moderates the effects of stressful life events on internalizing symptoms in children at familial risk for depression. Psychophysiology 49, 510–21 (2012).

[43] Mennella, R., Benvenuti, S.M., Buodo, G. & Palomba, D. Emotional modulation of alpha asymmetry in dysphoria: results from an emotional imagery task. International Journal of Psychophysiology 97, 113–119 (2015).

[44] Allen, J.J.B. & Reznik, S.J. Frontal eeg asymmetry as a promising marker of depression vulnerability: Summary and methodological considerations. Curr Opin Psychol 4, 93–97 (2015).

[45] Fava, G.A., Tomba, E. & Bech, P. Clinical pharmacopsychology: Conceptual foundations and emerging tasks. Psychotherapy and Psychosomatics 86, 134–140 (2017).

[46] Wakefield, J.C. & Schmitz, M.F. When does depression become a disorder? using recurrence rates to evaluate the validity of proposed changes in major depression diagnostic thresholds. World Psychiatry 12, 44–52 (2013).

[47] Berwian, I.M., Wenzel, J., Kasper, L., Veer, I., et al. Longer-term antidepressant use might normalise aberrant resting-state functional connectivity in remitted depression. Submitted (2019).

[48] Berwian, I.M., Wenzel, J., Collins, A.G., Seifritz, E., Stephan, K.E. et al. Computational mechanisms of effort and reward decisions in depression and their relationship to relapse after antidepressant discontinuation. Revision. under review, JAMA Psychiatry (2019).

[49] Wittchen, H.U. & Fydrich, T. Strukturiertes Klinisches Interview für DSM IV. Manual zum SKID-I und SKID-II. Hofgrefe, Göttingen (1997).

[50] Williams, J.B. A structured interview guide for the hamilton depression rating scale. Arch Gen Psychiatry 45, 742–7 (1988).

[51] Rush, A.J., Gullion, C.M., Basco, M.R., Jarrett, R.B. & Trivedi, M.H. The inventory of depressive symptomatology (ids): psychometric properties. Psychol Med 26, 477–86 (1996).

[52] Spitzer, R.L., Kroenke, K., Williams, J.B.W. & Löwe, B. A brief measure for assessing generalized anxiety disorder: the gad-7. Archives of internal medicine 166, 1092–1097 (2006).

[53] Treynor, W., Gonzalez, R. & Nolen-Hoeksema, S. Rumination reconsidered: A psychometric analysis. Cognitive Therapy and Research 27, 247–259 (2003).

[54] Huffziger, S. & Kühner, C. Die ruminationsfacetten broodung und reflection: Eine psychometrische evaluation der deutschsprachigen version rsq-10d. Zeitschrift für Klinische Psychologie und Psychotherapie 41, 38–46 (2012).

[55] Lehr, S. Mehrfachwahl-Wortschatz-Intelligenztest MWT-B. Spitta, Balingen (2005).

[56] Wechsler, D. Wechsler Adult Intelligence Scale-fourth edition (WAIS-IV). Psychological Corporation, San Antonio, Texas (2014).

[57] Reitan, R.M. Validity of the trial making test as an indicator of organic brain damage. Perceptual and Motor Skills 8, 271–276 (1958).

[58] Löwe, B., Decker, O., Müller, S., Brähler, E., Schellberg, D. et al. Validation and standardization of the generalized anxiety disorder screener (GAD-7) in the general population. Medical Care 46, 266–274 (2008).

[59] Winkler, I., Haufe, S. & Tangermann, M. Automatic classification of artifactual ica-components for artifact removal in eeg signals. Behavioral and brain functions 7, 1–15 (2011).

[60] Kayser, J. & Tenke, C.E. Principal components analysis of laplacian waveforms as a generic method for identifying erp generator patterns: I. evaluation with auditory oddball tasks. Clinical neurophysiology : official journal of the International Federation of Clinical Neurophysiology 117, 348–368 (2006).

[61] Smith, E.E., Reznik, S.J., Stewart, J.L. & Allen, J.J.B. Assessing and conceptualizing frontal eeg asymmetry: An updated primer on recording, processing, analyzing, and interpreting frontal alpha asymmetry. International journal of psychophysiology : official journal of the International Organization of Psychophysiology 111, 98–114 (2017).

[62] Grünewald, B.D., Greimel, E., Trinkl, M., Bartling, J., Großheinrich, N. et al. Resting frontal eeg asymmetry patterns in adolescents with and without major depression. Biological psychology 132, 212–216 (2018).

[63] Kołodziej, A., Magnuski, M., Ruban, A. & Brzezicka, A. No relationship between frontal alpha asymmetry and depressive disorders in a multiverse analysis of five studies. eLife 10, e60595 (2021).

[64] Segrave, R.A., Cooper, N.R., Thomson, R.H., Croft, R.J., Sheppard, D.M. et al. Individualized alpha activity and frontal asymmetry in major depression. Clinical EEG and neuroscience 42, 45–52 (2011).

[65] Deldin, P.J. & Chiu, P. Cognitive restructuring and EEG in major depression. Biological Psychology 70, 141–151 (2005).

[66] Rottenberg, J. & Hindash, A.C. Emerging evidence for emotion context insensitivity in depression. Current Opinion in Psychology 4, 1–5 (2015). Depression.

[67] Bylsma, L.M. Emotion context insensitivity in depression: Toward an integrated and contextualized approach. Psychophysiology 58, e13715 (2021).

[68] Stange, J.P., MacNamara, A., Barnas, O., Kennedy, A.E., Hajcak, G. et al. Neural markers of attention to aversive pictures predict response to cognitive behavioral therapy in anxiety and depression. Biol Psychol 123, 269–277 (2017).

[69] Bauer, E.A., Wilson, K.A., Phan, K.L., Shankman, S.A. & MacNamara, A. A neurobiological profile underlying comorbidity load and prospective increases in dysphoria in a focal fear sample. Biol Psychiatry 93, 352–361 (2023).

[70] Kanske, P., Heissler, J., Schönfelder, S. & Wessa, M. Neural correlates of emotion regulation deficits in remitted depression: the influence of regulation strategy, habitual regulation use, and emotional valence. Neuroimage 61, 686–693 (2012).

[71] Wade, E.C. & Iosifescu, D.V. Using electroencephalography for treatment guidance in major depressive disorder. Biological Psychiatry: Cognitive Neuroscience and Neuroimaging 1, 411–422 (2016).

[72] Widge, A.S., Bilge, M.T., Montana, R., Chang, W., Rodriguez, C.I. et al. Electroencephalographic biomarkers for treatment response prediction in major depressive illness: A meta-analysis. Am J Psychiatry 176, 44–56 (2019).

[73] American Psychiatric Association. Diagnostic and Statistical Manual of Mental Disorders. American Psychiatric Association Press (1994).

